# Serotonergic modulation of motor subspace dynamics drives a sleep-independent quiescent state

**DOI:** 10.64898/2026.01.28.702359

**Authors:** Kexin Qi, Yuming Chai, Guodong Tan, Daguang Li, Quan Wen

## Abstract

The dorsal raphe nucleus (DRN) serotonergic (5-HT) system has been implicated in regulating sleep and motor control; however, its specific role remains controversial. In this study, we found that optogenetic activation of DRN 5-HT neurons in larval zebrafish induced a quiescent state and a reduced response to acoustic stimuli. Unlike sleep, the induced quiescent state was not accompanied by a loss of postural control, and nighttime activation of DRN 5-HT neurons led to a subsequent sleep rebound. Whole brain light field imaging combined with demixed principal component analysis (dPCA) revealed distinct neural subspaces related to DRN activation, sound responses, and motor activity. DRN 5-HT activation selectively modulated the motor-related subspace while leaving the sound-evoked subspace unaffected. Unlike DRN activation, sleep induced by mepyramine significantly altered sound-evoked neuronal activity patterns. Further analysis demonstrated that serotonin had a graded effect on the motor subspace, wherein downstream neurons responsible for particular bout types were more significantly influenced. Embedding motor population activity in a curved geometric space revealed that the degree of curvature scales with behavioral suppression across animals, providing a quantitative signature of the quiescent state. Together, these results elucidate that serotonergic modulation promotes behavioral quiescence through selective regulation of motor populations.

## Introduction

Serotonin (5-hydroxytryptamine, 5-HT), known as the “feel-good” molecule, is a key monoaminergic neurotransmitter that exerts an expansive influence across the central nervous system (1–6), regulating essential physiological and behavioral processes such as locomotion (7–15), sleep (16–20), emotion (21–23), and learning (24–28). Within the vertebrate brain, the dorsal raphe nucleus (DRN) serves as the primary hub for serotonergic synthesis, providing the dominant source of ascending serotonin to widespread brain regions (2, 29). The DRN 5-HT system has long been implicated in the regulation of sleep and arousal; yet, its precise contribution to the sleep–wake cycle remains one of the most enduring enigmas in neurobiology (19, 30, 31). For more than half a century, a central controversy has persisted: whether serotonergic signaling functions as a primary driver of sleep induction or as a promoter of cortical arousal (16, 17, 19, 30–34).

The larval zebrafish (*Danio rerio*) provide an excellent model for reconciling these conflicting perspectives. They exhibit well-defined sleep behavior, characterized by reduced locomotor activity, sustained immobility, elevated arousal thresholds, and homeostatic sleep rebound following sleep deprivation (35–41). The zebrafish serotonergic system is highly conserved with mammals in anatomy and function (18, 42–44). The small size and optical transparency of the larval zebrafish brain enable whole-brain, cellular-resolution imaging, offering a unique opportunity to dissect serotonergic modulation across the entire neural population.

In zebrafish, increased DRN 5-HT neuron activity is often associated with locomotor suppression, suggesting that serotonergic signaling promotes sleep. Optogenetic activation of these neurons induces sleep-like behavior, while their ablation disrupts sleep-promoting pathways(18, 20), supporting a sleep-promoting role. Yet several lines of evidence suggest the opposite. Electrophysiological recordings show that DRN 5-HT neurons fire at higher rates during the day than at night (18), which is the opposite of the sleep rhythm, indicating a wake-related component of serotonergic activity. Transient water-flow stimulation and exposure to conspecific alarm substances both enhance DRN 5-HT neuron activity and sensory responsiveness, even when locomotion is suppressed (44, 45).

Beyond arousal and sleep, DRN 5-HT neurons are consistently more active in behavioral states with reduced locomotion. During exploitation in naturalistic foraging, zebrafish exhibit decreased movement, increased predation, and elevated DRN activity (46). DRN 5-HT neurons are essential for motor adaptation under visuomotor mismatch, where higher 5-HT levels correlate with reduced tail movements and reticulospinal excitability (47–50). These findings indicate that serotonergic activity is tightly linked to locomotor suppression across diverse behavioral contexts.

These seemingly contradictory studies pose a central question: How do changes in serotonin levels modulate an animal’s internal state and behavior, such that they either produce enhanced sensitivity to external stimuli, as in vigilance, or reduced sensitivity, as in sleep? We address this using a custom all-optical system for simultaneous optogenetic manipulation of DRN 5-HT neurons and whole-brain calcium imaging in behaving larval zebrafish. Zebrafish spontaneously alternate between locomotor and quiescent states, with DRN activity markedly elevated during quiescence. Optogenetic activation of DRN 5-HT neurons induced behavioral quiescence and reduced responsiveness to acoustic stimuli, in a state distinct from natural sleep. Whole-brain light-field imaging with demixed principal component analysis (dPCA) revealed separate neural subspaces associated with DRN activation, auditory processing, and motor activity. Our findings reveal that serotonergic modulation promotes behavioral quiescence through hierarchical control of motor circuits without altering auditory stimulus encoding, elucidating how the 5-HT system shapes brain states and behavioral flexibility.

## Results

### DRN 5-HT activation creates a quiescent, non-sleep state

We developed a custom optical system (51) for stable, long-term whole-brain imaging, tracking, and targeted optogenetic stimulation in freely swimming larval zebrafish (Fig. 1A). We recorded 60-minute spontaneous behavior and brain-wide neural activity in *elavl3:H2B-jGCaMP8s* transgenic larvae. The fish alternated naturally between locomotor and quiescent states (Fig. 1B). Consistent with previous work (46), whole-brain imaging revealed sustained and elevated DRN neuronal activity during the quiescent state (Fig. 1C, Fig. 1D). We then optogenetically activated DRN serotonergic neurons in the *Tg(tph2:ChrimsonR-mKate2)* transgenic fish (Fig. 2A). Although the *tph2* promoter primarily drives expression in DRN 5-HT neurons, minor expression in the pineal gland of the forebrain was minimized by spatially restricting 588 nm laser stimulation to the DRN (30 *µ*W, 50 Hz galvo scanning, Fig. 1A). Each 5-min stimulation was repeated 2–4 times per fish. DRN activation markedly suppressed locomotor activity, inducing a near-quiescent state, whereas control fish lacking ChrimsonR showed no effect (Fig. 2B). Low-frequency optogenetic stimulation also produced a similar reduction in locomotor activity (Fig. S1A-B).

**Figure 1.**
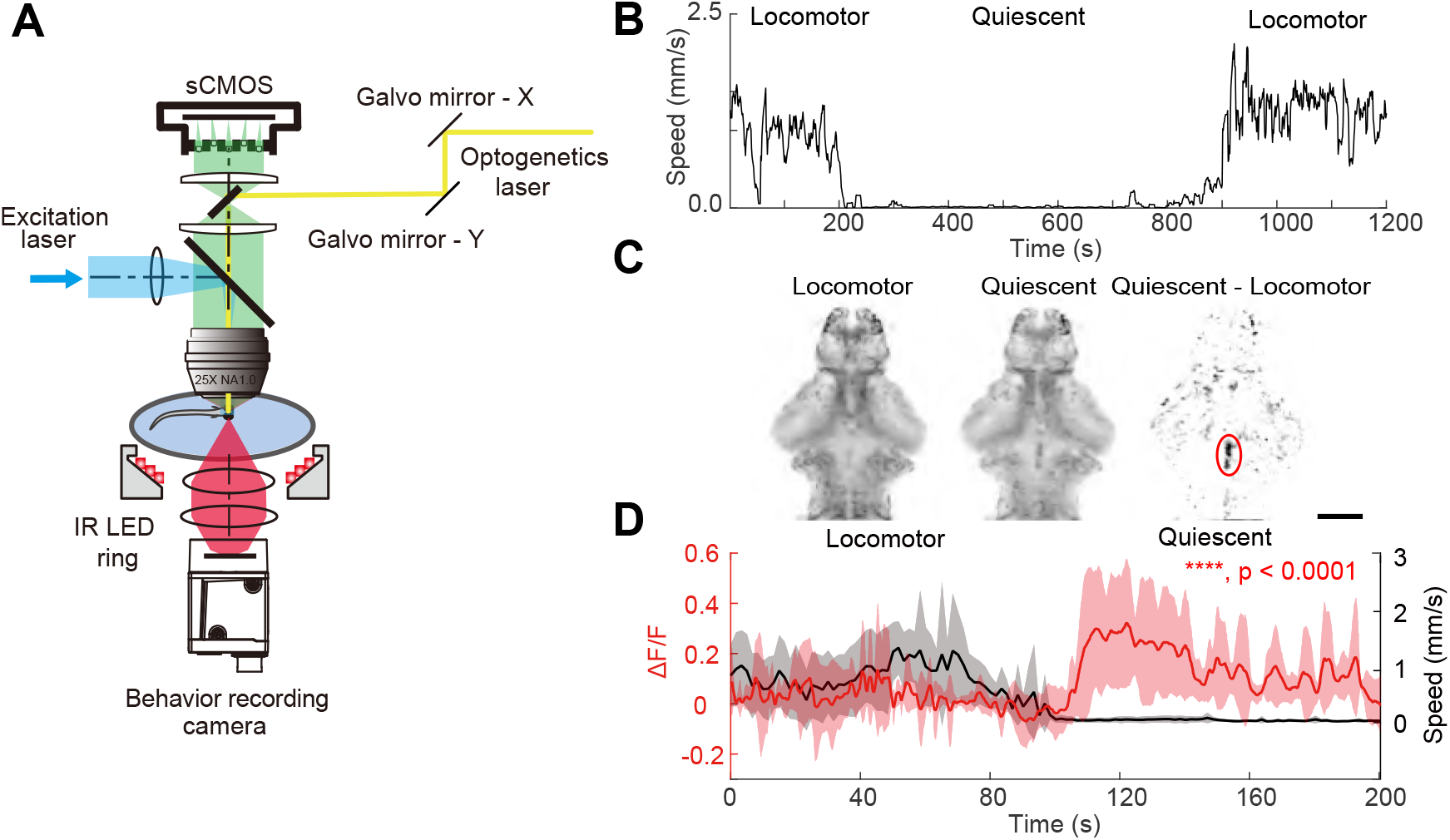
DRN neuronal activity increased during quiescence. **A**. A schematic of our all-optical system that integrates tracking, dual-color volumetric fluorescence imaging, and optogenetic manipulation. **B**. Example zebrafish exhibited alternating locomotor and quiescent states during spontaneous behavior. **C**. Maximum intensity projection (MIP) of whole-brain imaging in a zebrafish, showing 30 s averaged neural activity during locomotor (left) and quiescent (middle) states; their difference is shown on the right. The red circle marks the dorsal raphe nucleus (DRN). Scale bar, 100 *µ*m. **D**. Relationship between locomotor speed and DRN neural activity. Data are mean ± SD, n = 5 fish, p < 0.0001 (Mann–Whitney U test).

**Figure 2.**
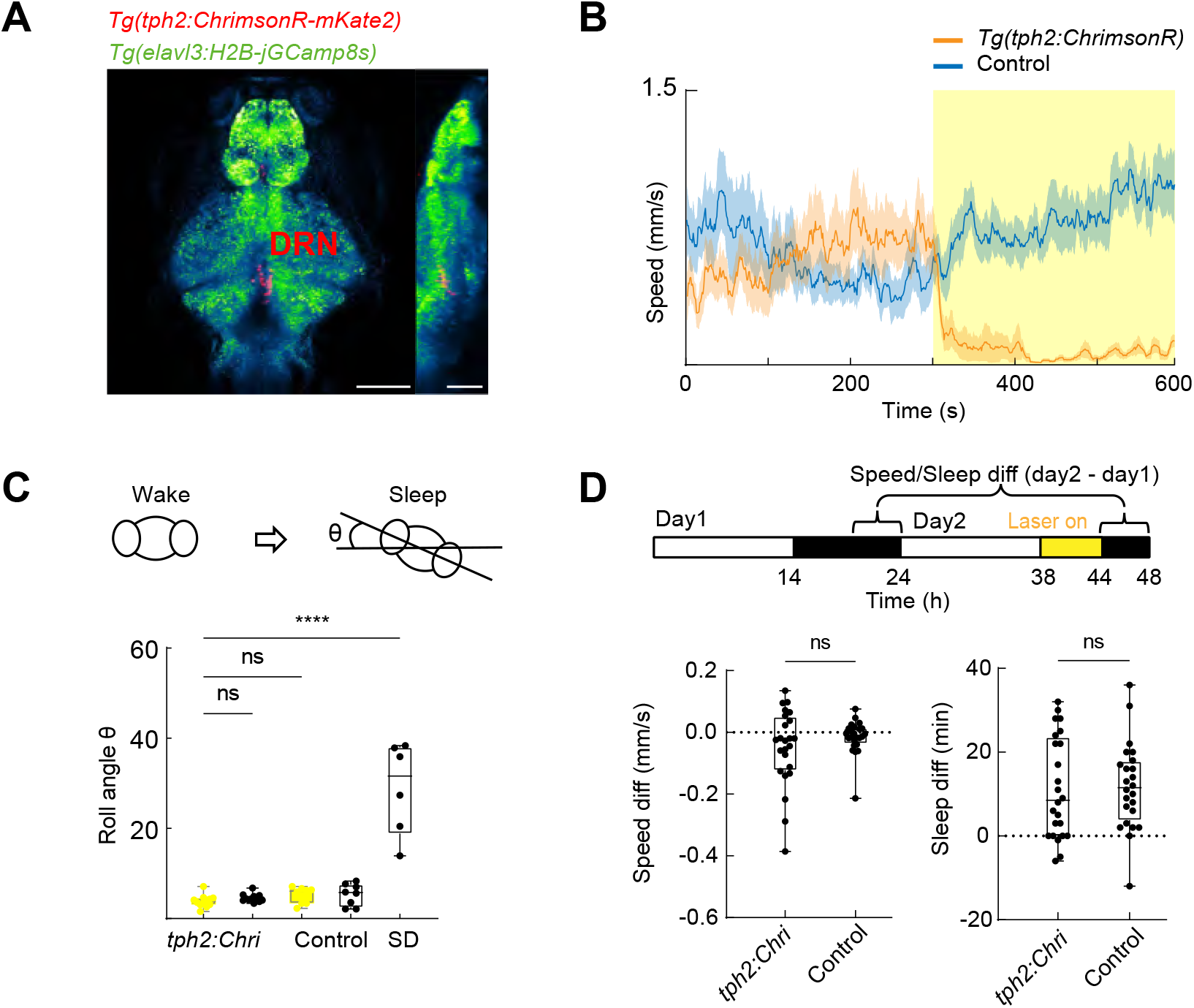
DRN 5-HT activation induces a quiescent but non-sleep-like state. **A**. MIP of whole-brain data from a 7 dpf *Tg(tph2:ChrimsonR-mKate2 × elavl3:h2b-jGCaMP8s)* zebrafish acquired by two-photon microscopy. Scale bar, 100 *µ*m. **B**. Locomotor velocity changes in *Tg(tph2:ChrimsonR)* and control zebrafish during DRN 5-HT neuron activation (n = 6). Yellow shading marks optogenetic stimulation. **C**. Top: Body roll angle (rotation in the Y–Z plane) increases during natural sleep, indicating loss of postural stability. Body roll angle in control (n = 8), sleep-deprived (SD, n = 6), and *Tg(tph2:ChrimsonR)* zebrafish (n = 12). Yellow indicates optogenetic stimulation. In *Tg(tph2:ChrimsonR)* fish, light versus no-light conditions were compared with the Wilcoxon matched-pairs signed rank test. *Tg(tph2:ChrimsonR)* versus control and sleep-deprived versus *Tg(tph2:ChrimsonR)* were compared with the Mann–Whitney U test. ****p < 0.0001. **D**. Top: Experimental timeline over two light–dark cycles (14 h light/10 h dark). The first cycle was normal; in the second, optogenetic stimulation was applied during the first 6 h of the dark period. Average locomotor speed and sleep duration in the 4 h after stimulation were compared with the corresponding 4 h of the first dark period. Bottom: Differences in average locomotor speed and sleep duration between *Tg(tph2:ChrimsonR)* (n = 24) and control zebrafish (n = 24), analyzed with the Mann–Whitney U test.

Optogenetic activation of DRN 5-HT neurons suppressed locomotion in freely swimming zebrafish. This effect has been interpreted as either promoting sleep (18) or vigilance-related immobility (44). To test whether this quiescence resembles sleep, we examined postural control. Unlike natural sleep, which causes postural instability and increased roll angles(52), DRN 5-HT activation did not significantly alter roll angles, whereas sleep-deprived (SD) zebrafish showed clear instability during quiescence (Fig. 2C, Fig. S1C-D). We next tested whether DRN activation during the dark (sleep) phase would block sleep rebound: if this quiescent state were sleep, no rebound would be expected. Instead, stimulation led to a rebound comparable to controls (Fig. 2D, Fig. S1E-F). Together, these results suggest that DRN activation induces a quiescent but non-sleep state in larval zebrafish.

### DRN 5-HT activation modulates brain state

To assess how optogenetic activation of DRN 5-HT neurons alters global neural dynamics, we applied demixed principal component analysis (dPCA) to whole-brain activity data (53). dPCA decomposes population activity into components associated with specific experimental variables, enabling the isolation of DRN-dependent effects. To minimize direct optogenetic effects on the identified subspace, DRN-localized regions were excluded from the analysis (Methods).

Within this analysis framework, each dPCA component can be interpreted as a direction in population activity space along which neural activity varies with a given variable (e.g., optogenetic activation). Projecting population activity onto this axis yields a low-dimensional trajectory that captures how this variable modulates neural dynamics over time (Fig. S2). The strength of modulation can be quantified as the fraction of variance associated with the variable that is explained by the components, and neurons with larger weights contribute more strongly to this variable-related activity.

Components associated with DRN activation explained significantly more variance in *Tg(tph2:ChrimsonR)* zebrafish than in controls (Fig. 3A), indicating a strong serotonergic impact on brain-wide neural activity. The small stimulation-related variance in controls likely reflected visual responses to the laser. Projection of whole-brain activity onto the first demixed principal component (dPC1 score), which accounted for over half of the data variance (Fig. 3A), revealed a pronounced and reversible state transition during DRN activation (Fig. 3B). Brain regions with high dPC1 weights (Fig. 3C) showed activity more strongly correlated with locomotor behavior than randomly selected regions (Fig. 3D and Method). Thus, DRN 5-HT activation rapidly reorganizes global neural states and selectively engages motor-related circuits.

**Figure 3.**
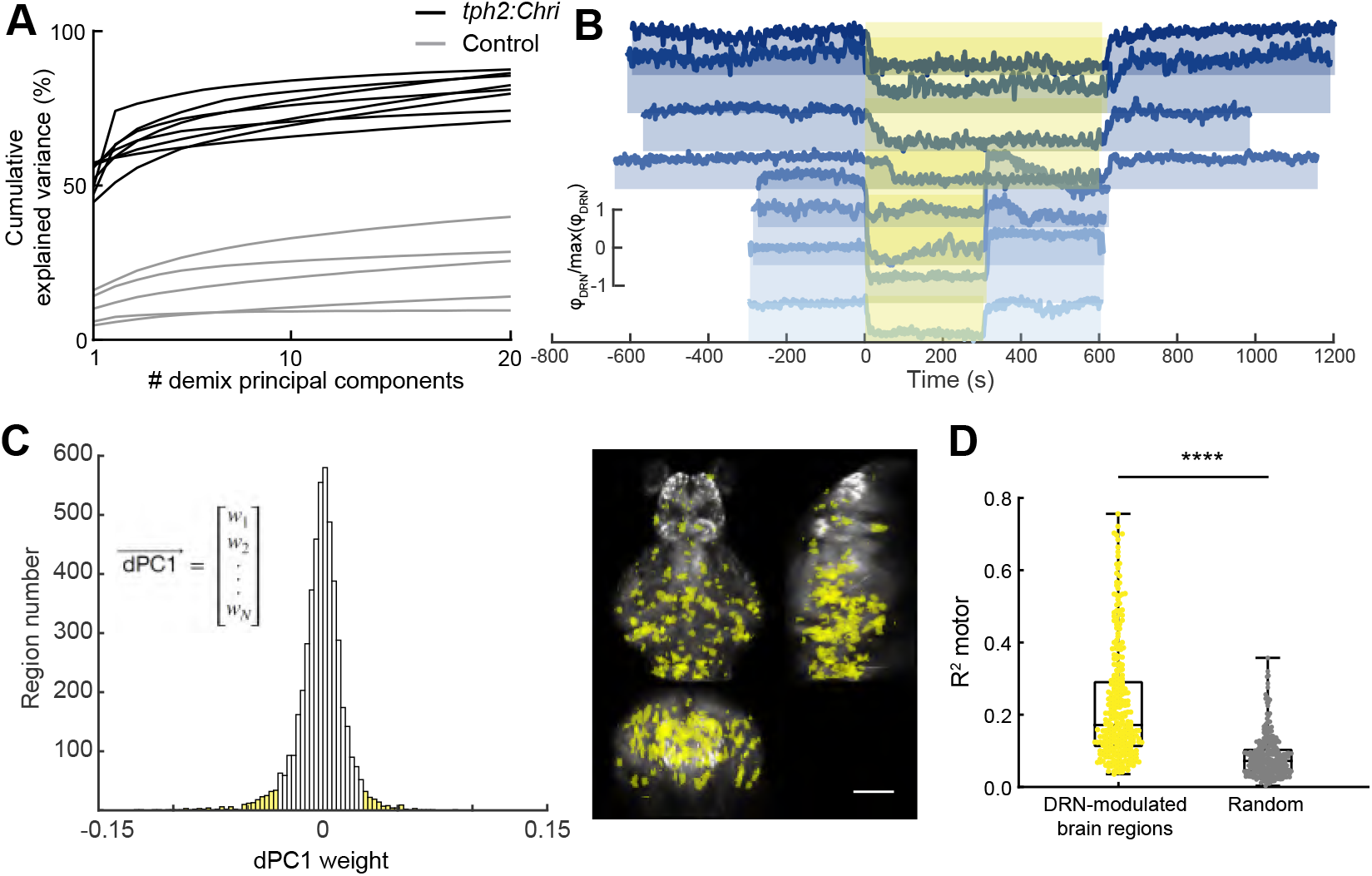
DRN 5-HT activation modulates brain state. **A**. Cumulative variance explained by demixed principal components (dPCs) related to optogenetic stimulation in *Tg(tph2:ChrimsonR)* zebrafish (n = 8) and controls (n = 5). **B**. Time course of whole-brain activity projected onto dPC1 in *Tg(tph2:ChrimsonR)* zebrafish (n = 8). Yellow shading marks optogenetic stimulation. *ϕDRN* represents dPC1 score. **C**. Left: Histogram of brain-region weight distribution in dPC1. Yellow shading highlights high-weight regions (|weight| *>* 0.03, 272 regions). Right: Spatial distribution of these regions in the zebrafish brain. Scale bar, 100 *µ*m. **D**. *R*^2^ between neural activity in dPC1 high-weight regions and locomotor behavior, compared with randomly selected regions (n = 272; Mann–Whitney U test, ****p<0.0001).

### DRN 5-HT activation modulates motor circuits to suppress sound-evoked responses

Across species, sleep features an increased arousal threshold and reduced responsiveness to external stimuli (54). In contrast, a recent study shows that DRN 5-HT neuron activation enhances zebrafish vigilance to aversive cues (44), lowering the auditory response threshold – an effect opposite to the sleep-promoting effect of DRN activation (18). To test how DRN 5-HT neurons regulate behavioral states, we present auditory stimuli during DRN activation. Based on larval auditory sensitivity (55), a 350 Hz pure tone (500 ms, 100 dB) was delivered every 60 s (Fig. 4A). *Tg(tph2:ChrimsonR)* larvae exhibited strong locomotor suppression and near-complete loss of sound-evoked escape responses during stimulation (Wilcoxon matched-pairs signed rank test, **p = 0.0039), both of which recovered afterwards (Fig. 4B). Control fish showed no significant changes in swimming speed or escape probability but displayed weak habituation across trials (Fig. 4B, (56)).

**Figure 4.**
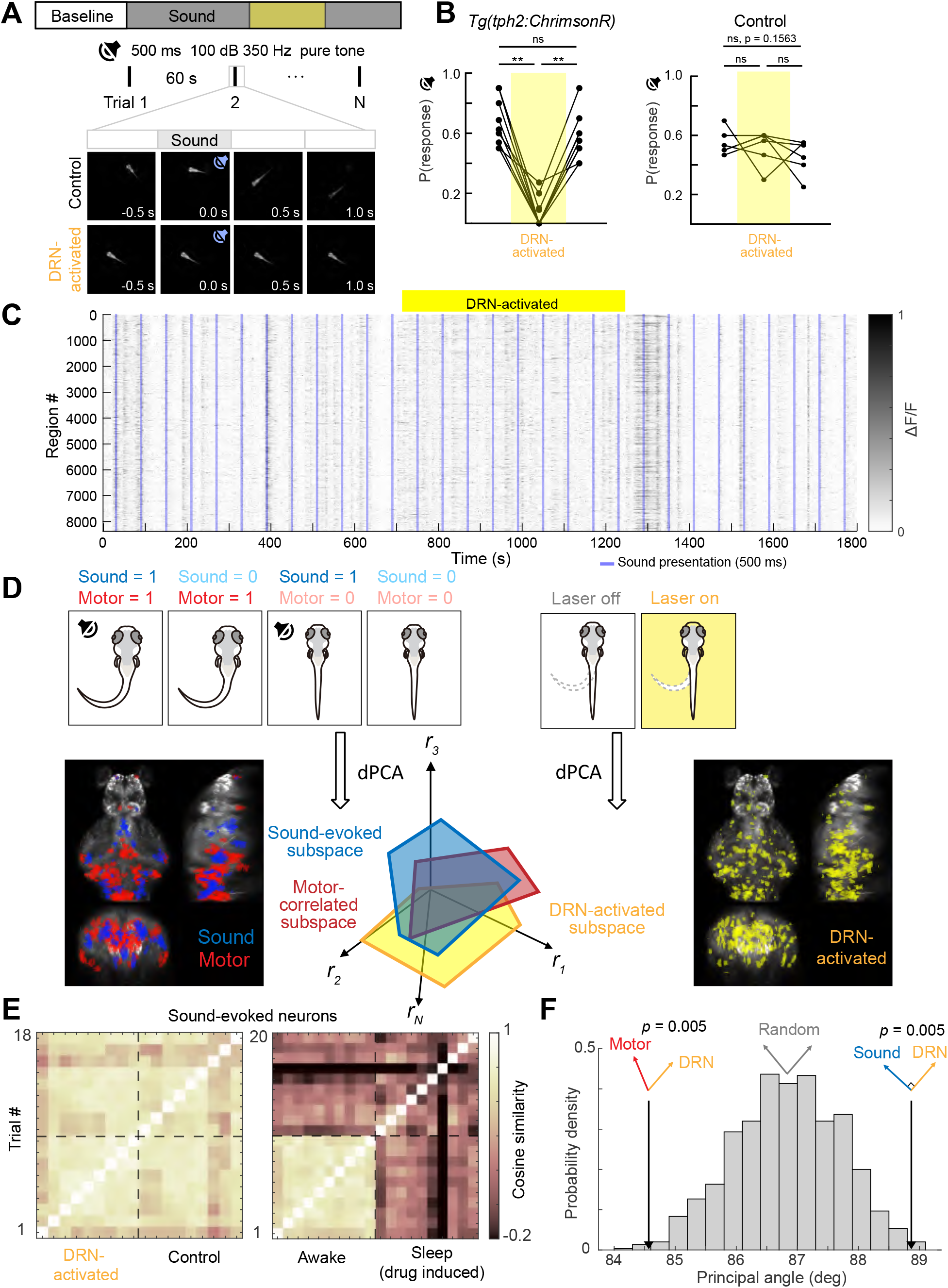
DRN 5-HT activation modulates motor circuits to reduce sound-evoked responses. **A**. Experimental protocol for sound stimulus experiments. **B**. Probability of sound-evoked escape in *Tg(tph2:ChrimsonR)* and control zebrafish before, during, and after optogenetic stimulation. Yellow shading marks optogenetic activation. Wilcoxon matched-pairs signed rank test, **p = 0.0039. **C**. Population raster plot of simultaneously recorded neurons during DRN 5-HT activation and control. Blue lines mark sound onset. **D**. Schematic of sound, motor, and DRN activation subspaces identified by dPCA. Left and right MIPs show brain regions with high weights in each subspace in an example zebrafish (bottom right panel is the same as the panel shown in Figure 3C). **E**. Left: Similarity matrix of sound-evoked population responses during DRN activation vs. control. Right: Same analysis comparing awake and drug-induced sleep. **F**. Principal angle analysis shows the motor subspace is significantly aligned with the DRN activation subspace, while the sound subspace is nearly orthogonal (p-values from a nonparametric permutation test, 1000 iterations). Arrows show the mean angle, across all fish, between the DRN 5-HT activation subspace and the motor-related subspace (left) or the sound-related subspace (right). “Significantly aligned” means the motor–DRN angle is significantly smaller than the random baseline (gray), and “significantly orthogonal” for sound–DRN means the angle is significantly closer to 90^◦^ than the random baseline.

To examine how DRN 5-HT neuron activation affects sensorimotor processing (Fig. 4C), we next recorded whole-brain neural activity in head-fixed, tail-free larvae embedded in agarose to capture transient calcium signals with minimal motion artifacts. dPCA revealed neural subspaces associated with auditory processing, motor activity, and DRN 5-HT activation (Fig. 4D, Fig. S3A). We quantified trial-to-trial similarity of sound-evoked subspace activity using cosine similarity, which compares the angle between population activity vectors rather than their magnitude.

Values closer to 1 indicate more similar activity patterns across trials. Interestingly, DRN activation preserved the structure of the auditory population code (Fig. 4E, Fig. S3B). Because head-fixed larvae rarely enter natural sleep, we applied 1 mM mepyramine, a sleep-promoting antihistamine, to induce a sleep-like state (41), which markedly changed auditory responses (Fig. 4E, Fig. S3C). Using both strong and weak auditory stimuli (Fig. S3D), we found no evidence that DRN 5-HT activation altered sound-evoked responses at either intensity (Fig. S3E). Compared against a null distribution – the expected principal angle between two random activity subspaces – our analysis revealed a statistically significant alignment between DRN activation and motor-related neural subspaces, with the sound-related subspace being nearly orthogonal (Fig. 4F and Methods). Here, alignment refers to the geometric relationship between high-dimensional neural subspaces rather than overlap of individual neurons. Thus, DRN 5-HT neuron activation selectively modulates motor-related activity while preserving auditory encoding, thereby reshaping sensorimotor processing.

### DRN 5-HT neuron activation produces bout type–dependent, graded suppression of the motor subspace

We constructed a linear regression model using baseline tail movements to predict neural activity (57). After detecting bouts, we computed each bout’s direction and amplitude and classified them into 12 types (Fig. S4). Based on the timing of each bout type, we defined 12 regressors (*r*_1_–*r*_12_) with corresponding coefficients (*β*_1_-*β*_12_) (Fig. 5A, Methods). Using spontaneous behavioral data and motor-correlated dPC1 weight (Fig. S5A), we identified motor-correlated neurons and quantified their coding selectivity using the coefficient of variation (CV) of their 12 regression coefficients. Some neurons showed similar activity across all bout types, yielding low coefficient variability (Fig. 5B top); whereas others responded selectively to specific bout types – such as those involving larger tail amplitudes and turning angles (for instance, type 12) – and therefore displayed higher variability in regression coefficients (Fig. 5B bottom).

**Figure 5.**
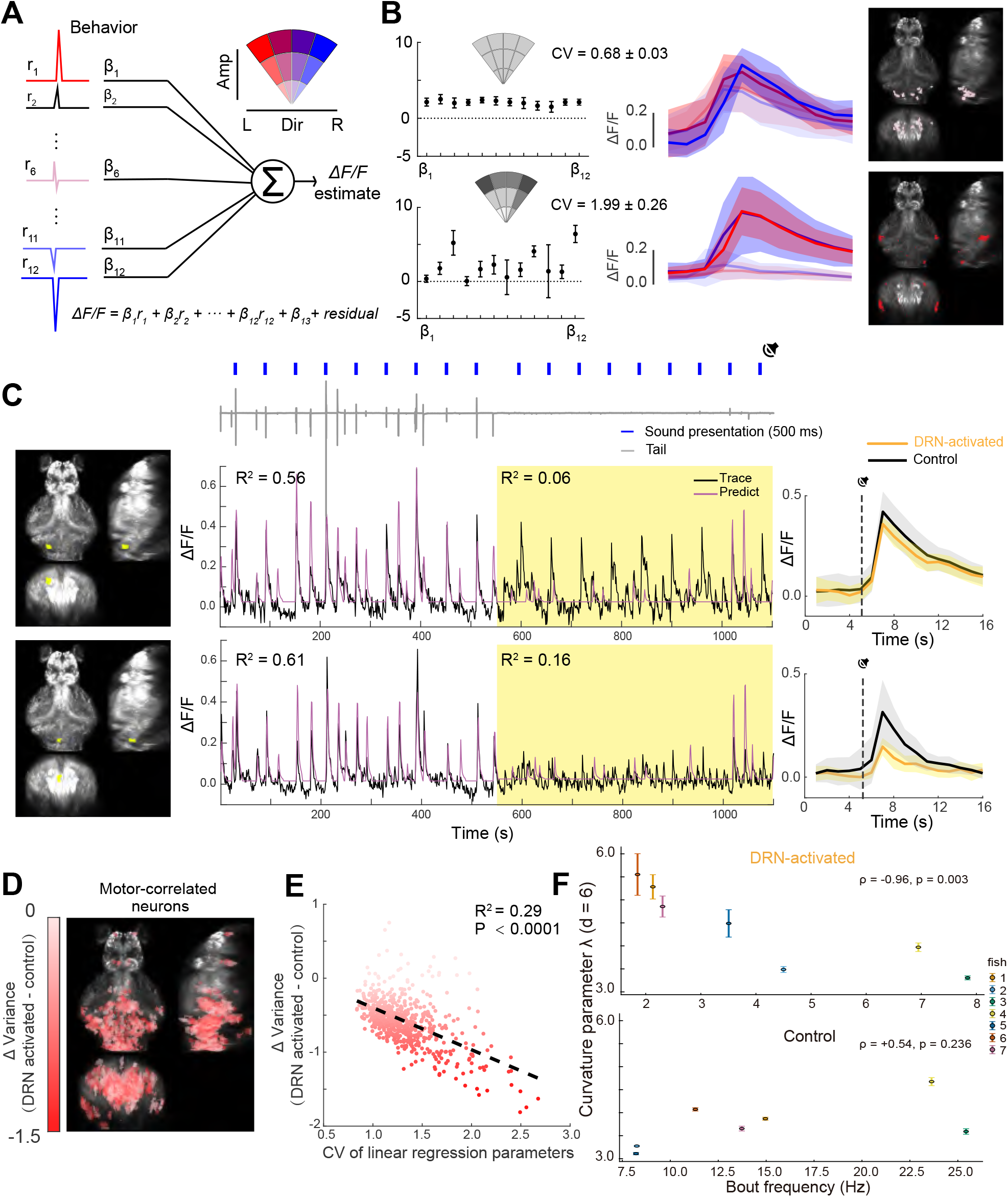
DRN 5-HT neuron activation exerts graded suppression on motor subspace. **A**. Schematic of the linear regression analysis. **B**. Example neurons with low (top) and high (bottom) variability in regression coefficients. Left, middle, and right panels show regression coefficients, mean activity across bout types, and neuronal spatial locations. **C**. Two motor-related regions with distinct modulation after DRN 5-HT activation. Left, middle, and right panels show their spatial locations, activity in control and optogenetic trials, and trial-averaged activity. **D**. Spatial distribution of motor-correlated neurons differentially modulated by DRN 5-HT activation. Neural magnitude is quantified by variance. Neurons are color-coded by variance reduction (darker red, stronger suppression; lighter red, weaker effect), as in (Fig. 5E,F). **E**. Relationship between DRN 5-HT–induced modulation and the coefficient of variation (CV) of regression coefficients across motor-correlated neurons. **F**. Hyperbolic curvature of the motor population tracks behavioral quiescence during DRN 5-HT activation. Each point is one fish (n = 7), colored by individual. Y-axis: curvature parameter *λ* from Bayesian hyperbolic MDS of pairwise motor-population correlation distances (*d* = 6; *K* = *λ*^2^); error bars, 1 SD. X-axis: bout frequency in the same period. Top, optogenetic laser on: stronger behavioral suppression corresponds to higher curvature (Spearman *ρ* = 0.96, *p* = 0.003, exact permutation test; Pearson *r* = 0.87, *p* = 0.011). Bottom, laser off: no significant relationship in baseline (*ρ* = +0.54, *p* = 0.236, Pearson *r* = +0.59, *p* = 0.166).

We next applied the linear model to predict neural activity during both control and DRN 5-HT activation periods. Although zebrafish exhibited markedly reduced locomotion during DRN 5-HT activation, some motor-correlated neurons maintained control-like activity levels (Fig. 5C, top). However, the neural activity no longer drove movements, indicating a decoupling between neuronal activity and behavior and reducing the variance in neural activity explained by the linear model. In contrast, other neurons displayed a pronounced reduction in activity during 5-HT activation (Fig. 5C, bottom). Using the differences in neuronal activity between control and DRN 5-HT activation, we mapped the spatial distribution of motor-correlated neurons with distinct modulation patterns (Fig. 5D). Interestingly, neurons least affected by DRN 5-HT activation (light red) had lower variability in their regression coefficients, whereas the most affected neurons (dark red) had higher variability (Fig. 5E, Fig. S4B-D). Together, these results suggest that serotonergic modulation exerts a graded suppression on the motor network, preferentially inhibiting downstream neurons involved in specific motor actions to reduce locomotion.

We next asked how DRN 5-HT activation reshapes the geometry of motor-related population activity. We applied Bayesian multi-dimensional scaling (MDS) to the matrix of pairwise neural correlation distances (58) (Fig. S7A). For each neuron pair (*i, j*) with Pearson correlation *C*_*ij*_, we defined 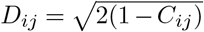 (Methods) so that perfectly correlated neurons have a distance of 0, uncorrelated neurons have a distance of 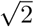, and perfectly anti-correlated neurons have a distance of 2. MDS then finds a low-dimensional arrangement of points that best preserves these distances – nearby points have similar activity, and distant points are dissimilar – yielding a geometric snapshot of the population’s functional organization. In the Bayesian framework, the geometry of this space is not fixed *a priori*: it compares candidate embeddings and selects the one that best preserves the distances. We tested both flat Euclidean space and hyperbolic space. Hyperbolic space expands exponentially from any point, so distances near its boundary grow much faster than in a Euclidean space of the same dimension (Fig. S6), making it well suited to data with a hierarchical structure (59, 60), or, more generally, to data in which subsets of points become strongly separated from the rest.

For each fish, we fitted separate embeddings for DRN activation (laser on) and baseline (laser off). The optimal embedding dimension *d* was between 6 and 8 (Bayesian Information Criterion, see Methods). Hyperbolic embeddings outperformed Euclidean ones in both conditions (Fig. S7C), with curvature parameters *λ* and *K* = − *λ*^2^ ranging from 3.3 to 5.6 across animals and experimental conditions. To test whether the hyperbolic fit captures a meaningful geometrical structure beyond the correlation spectrum, we randomized the eigenvectors (PCA weights) of the covariance matrix while preserving its eigenvalues (Methods). This surrogate still admitted a hyperbolic fit with large *λ*, but the fit showed systematic distortions: large embedding distances were compressed, deviating from the original distances (Fig. S9A). Thus, the quality of the hyperbolic embedding – its ability to preserve pairwise distances – depends on the specific neuronal arrangement in correlation space.

To further test whether this geometric measure reflects a behavior-dependent reorganization of the motor population rather than the idiosyncrasy of a single recording, we compared the fitted curvature to behavior across animals (Methods). For each fish, we used bout frequency during DRN activation as a readout of how strongly the network was driven toward quiescence (lower frequency = stronger suppression). Across fish (n = 7), laser-on curvature at a given embedding dimension closely tracked this measure (Fig. 5F, Fig. S8): fish in which DRN stimulation nearly abolished swimming showed the highest curvatures, whereas fish with weaker suppression showed lower curvatures (Spearman *ρ* = − 0.96, *p* = 0.003, exact permutation test). This coupling was specific to the activated state: during baseline in the same animals, curvature did not significantly relate to spontaneous bout frequency (*ρ* = +0.54, *p* = 0.236), and the trend reversed. Thus, motor-population embedding curvature is a state-dependent geometric signature that appears specifically when DRN 5-HT activation drives the network into quiescence, linking a single population-level parameter to a behavioral phenotype.

## Discussion

Activation of DRN 5-HT neurons generally suppresses locomotion in zebrafish, a phenomenon controversially interpreted as sleep or vigilance (18, 44). In this study, DRN 5-HT activation nearly abolished locomotion without altering body posture, and nighttime stimulation still produced normal sleep rebound. Thus, the induced quiescent state represents motor suppression rather than sleep. Unlike mammals, whose sleep states are determined through electroencephalogram (EEG) and electromyogram (EMG) recordings, zebrafish sleep is typically inferred based on several minutes of sustained immobility due to technical limitations (18, 36, 40). Since movement inhibition diminishes responsiveness to external stimuli, this resting yet awake state may be misclassified as sleep in zebrafish.

Indeed, multiple non-sleep behavioral states can also produce pronounced locomotor suppression; previous studies have demonstrated that serotonin contributes to movement suppression across diverse behavioral contexts (44, 46, 48, 49, 52, 61). In our study, zebrafish exhibited spontaneous alternations between locomotor and quiescent states, resembling the previously described “exploitation” and “exploration” modes (46). However, because no prey were present in our experiments, the observed locomotor reduction cannot be equated with the exploitation state. Elevated DRN serotonergic activity has been reported during both REM-like sleep (52) and quiet vigilance (44), and the latter is also associated with low-frequency synchronized forebrain activity (44). A common feature in these distinct states is the pronounced reduction in movement. We therefore propose that DRN serotonergic activation is not sufficient to define a specific behavioral state but primarily drives motor suppression. Consistent with this view, although DRN 5-HT activation strongly suppressed locomotion, we did not observe similar low-frequency synchronized forebrain dynamics during stimulation.

Single-cell connectivity analyses have shown that DRN 5-HT neurons project directly to midbrain and hindbrain regions involved in motor control (62). Consistent with this anatomical organization, whole-brain serotonin imaging has revealed elevated 5-HT levels in the hindbrain during states of reduced motor vigor, where inhibitory HTR1 receptors are predominant (49). At the circuit level, recent voltage imaging studies demonstrate that serotonergic neurons are dynamically regulated by local inhibitory inputs. Swim commands recruit GABAergic inhibition onto DRN 5-HT neurons, leading to a transient reduction in their activity during movement execution (50).

This serotonin-driven motor suppression function appears evolutionarily conserved. In *C. elegans*, activation of serotonergic neurons NSM induces robust locomotor suppression through specific receptor subtypes (10, 15). In mammals, although the serotonergic system has traditionally been regarded as part of the arousal-promoting network, numerous studies have also reported that elevated 5-HT levels suppress locomotion (9, 63–66). This kind of wakeful quiescent state serves important physiological functions. During intense or prolonged physical activity, serotonin levels in the brain and spinal cord rise, inhibiting motor neuron activity and inducing central fatigue, thereby reducing movement to prevent muscle and other bodily damage (67, 68). In a recent study, sick mice activated IL-1R1–expressing serotonergic neurons in the DRN via cytokine signaling, which in turn triggered social withdrawal and reduced locomotion—responses that promote recovery and limit pathogen transmission (66).

In our study, activating DRN 5-HT neurons did not significantly alter the representation of auditory information. The effects on other senses, such as vision or nociception, may differ, and the animal’s state could also modify how 5-HT influences sensory encoding. Previous research has observed 5-HT enhancing sensory information encoding under specific conditions (44, 45). In addition, previous studies have shown that under high-threat or punishment-based behavioral paradigms in mice (13), activation of DRN 5-HT neurons can instead increase locomotor activity. Consistently, the environmental threat level may also modulate the effects of DRN serotonergic activity on locomotion in zebrafish.

Finally, our analysis revealed that DRN 5-HT activation does not uniformly inhibit the motor network but instead exerts graded suppression. Neurons with broad tuning across multiple bout types remained largely unaffected, whereas those selectively responsive to specific motor patterns — particularly high-amplitude or turning movements — were strongly suppressed. We also show that the correlation structure of the motor population is well captured in a hyperbolic space, whose curvature serves as a geometric marker of quiescence. These results suggest that serotonergic modulation acts hierarchically at multiple levels of the motor circuitry, preferentially dampening specialized components that drive vigorous or directional movements. Such selective suppression may serve as a mechanism to reduce overall motor output while maintaining baseline motor tone, thereby facilitating rapid transitions between active and quiescent states. This implies that “no-action” is not merely the absence of movement but a strategic policy that prioritizes future state accessibility over immediate output (69), thereby creating the low-noise ‘offline’ environment (70) essential for learning, neural replay, and memory consolidation.

## ACKNOWLEDGEMENTS

The authors are grateful to Danqian Liu for experimental suggestions, Yu Mu for sharing the optogenetic fish line, Takashi Kawashima for sharing unpublished datasets, and Rubén Moreno Bote for discussions. This research was supported by the STI2030–Major Projects (Grant No. 2022ZD0211900), under the subproject “New Technologies for Whole-Brain-Scale Neuronal Mesoscopic Atlas” (Grant No. 2022ZD0211904).

## Methods

### Zebrafish

All larval zebrafish were raised in 0.5 × E2 Embryo Media at 28.5°C and a 14/10 hr light/dark cycle. The 0.5 × E2 Embryo Media consists of 7.5 mM NaCl, 0.25 mM KCl, 0.5 mM MgSO_4_, 75 µM KH_2_PO_4_, 25 µM Na_2_HPO_4_, 0.5 mM CaCl_2_, 0.35 mM NaHCO_3_. Larval zebrafish aged 6-11 days post-fertilization (dpf) were used for all experiments. Sex discrimination was not included since the sex of zebrafish is not specified at this stage.

### All-optical system

Neural activity and behavior were recorded using a custom-built all-optical system, as previously described (51). The system consists of three main components: a 3D tracking module, a dual-color fluorescence imaging module, and an optogenetic manipulation module. A convolutional neural network (CNN) was used to detect the lateral positions of the fish’s head from dark-field images captured by the near-infrared (NIR) tracking camera. A tracking model converted the real-time positional information into analog signals to drive the high-speed motorized stage and compensate for fish movement. An autofocus camera with a microlens array acquired multi-perspective images; the fish’s z-position was estimated via light-field principles, and a PID controller used this data to drive a piezo for axial tracking. The neural activity-dependent green fluorescence signal and the activity-independent red fluorescence signal were split into two beams by a dichroic mirror before entering the two sCMOS cameras separately. Both the red and green fluorophores can be excited by a blue laser (488 nm). A 2D galvo system deflected a yellow laser (588 nm) to a user-defined ROI in the fish brain for real-time optogenetic manipulation with the aid of a fast whole-brain image reconstruction and registration algorithm.

### Experimental procedures for whole-brain imaging and optogenetic manipulation

For freely swimming zebrafish, we employed a custom-designed circular chamber with a diameter of 20 mm and a height of 800 µm. The top and bottom surfaces of the chamber were coverslips, ensuring unobstructed optical access from both above and below. To prevent the chamber edges from occluding fluorescent signals, a 1.5% low-melting-point agarose solution (Thermo Fisher Scientific, product no. 16520050) was used to form an approximately 1 mm wide annular agarose ring around the chamber perimeter. For zebrafish not expressing optogenetic proteins ChrimsonR, the 488 nm laser was operated at 30 mW with a 10 Hz flashing mode and an exposure time of 1 ms per pulse. For zebrafish expressing optogenetic proteins, a long-exposure mode with a reduced power of 0.3 mW was applied to minimize potential interference with the ChrimsonR.

For head-fixed zebrafish, 1.5% low-melting-point agarose was used for immobilization. After the agarose had fully solidified, the agarose surrounding the head (for pharmacological experiments) and tail was carefully removed. Because head fixation substantially reduces motion-induced noise, we used a lower-power 488 nm excitation (0.03 mW) to further minimize interference with the optogenetic proteins ChrimsonR.

In the all-optical system, we incorporated a high-precision patterned optogenetic stimulation module that allowed selective activation of 5-HT neurons within the DRN in both freely swimming and head-fixed zebrafish. The stimulation pattern was spatially restricted to user-defined regions and avoided illumination of the pineal gland. Optogenetic activation was performed using a 588 nm yellow laser. A ROI (160 *×* 160*µ*m) was positioned to fully encompass the DRN. The laser power was 30 µW, and Galvo-based scanning produced an effective stimulation frequency of 50 Hz. Each optogenetic trial lasted 5 min, with 2–4 repetitions per fish. For experiments that included auditory stimulation, the duration of optogenetic activation was extended as needed to ensure sufficient trial numbers.

### Whole-brain image processing pipeline

Whole-brain neural activity was extracted from reconstructed 3D image stacks acquired by the all-optical system. Because both channels originate from the same optical path, spatial transformations are equivalent; registration was therefore first performed on the red channel, which is unaffected by neural activity, and the resulting transformation was applied to the green channel.

Registration included cropping, rigid registration, and non-rigid registration. Reconstructed 3D images (600 × 600 × 250 voxels) were rotated to a standard orientation based on head orientation and cropped to 308 × 380 × 210 voxels to match the ZBB standard brain template. A sharp reference frame without motion blur was selected; all other frames were aligned to this reference and subsequently registered to the standard brain using affine transformations from the Computational Morphology Toolkit (CMTK, NIH). After rigid registration, small deformations caused by physiological movements were corrected using Demons-based diffeomorphic non-rigid registration. To improve alignment accuracy, an averaged template was generated from 10 frames per 100 frame sequence, and each frame was aligned to this template using optical flow-based registration.

After applying the transformation from the red channel to the green channel, brain regions were segmented based on voxel-wise temporal correlations of the green channel fluorescence. For each voxel, the average correlation with its 14 neighboring voxels was computed to generate a correlation map. The map was segmented using a watershed algorithm to define preliminary regions, and voxels with low correlation to the regional mean were removed. The resulting segmentation was then applied to the red channel.

### Behavioral recording and sleep analysis

The experiment began at 18:00 on Day 1 and continued until 12:00 on Day 3, covering two dark cycles. To avoid edge obstruction during imaging, zebrafish larvae were housed in a custom-designed 24 well plate. The light/dark cycle was maintained using white LED illumination (lights on at 08:00 and off at 22:00). A 940 nm infrared light source was used for video recording. Optogenetic stimulation was delivered using a 530 nm LED light source (THORLABS M530L3) during the first 6 hours of the second dark phase. At 14:00 on Day 2, E2 embryo medium and live paramecia were added to prevent dehydration and starvation. Infrared videos were acquired at 1 fps using a Basler acA2000-165kmNIR camera. Illumination control and data acquisition were managed by a custom-written C++ program.

Videos were analyzed using ZebraZoom (https://zebrazoom.org/) to extract the locomotor activity of zebrafish larvae. Periods during which swimming speed remained below 0.0667 mm/s (corresponding to 1 pixel in the video, to avoid detection error) for at least 60 s were defined as a quiescent minute. To exclude abnormal data, episodes of continuous immobility lasting longer than 1 hour were removed. The average swimming speed and total immobility duration during the last 4 hours of the first dark phase were then calculated and compared with those from the last 4 hours of the second dark phase to assess the effects of optogenetic stimulation on locomotor activity and sleep-related behavior.

### Body roll angle

Zebrafish larvae were approximated as rigid bodies; therefore, the whole brain roll angle was used to estimate the body roll angle. Three-dimensional whole-brain fluorescence data were used for this calculation. Because only the roll angle was required and details of neural activity were not considered, images were downsampled from 2048 × 2048 to 512 × 512 pixels to accelerate processing, followed by deconvolution with 10 iterations. After reconstruction, the 3D images were binarized using adaptive thresholding, and the largest connected component was identified as the head region. Volumes with the largest connected component smaller than 100,000 voxels were considered to correspond to cases in which the fish deviated from the field of view; these data were excluded and filled by interpolation. Principal component analysis (PCA) was applied to determine its principal axis, representing the longitudinal axis of the brain. The brain was then rotated to align this axis with that of a standard reference brain. After alignment, a maximum intensity projection of the 3D image onto the Y–Z plane was generated, and PCA was applied to this projection to determine its principal axis. The angle between this axis and the horizontal vector is defined as the body roll angle of the zebrafish.

At 6 dpf, zebrafish reared under a normal light–dark cycle were subjected to continuous sleep deprivation for 3 days (41). Larvae were placed in a circular tank containing 200 mL of 0.5× E2 embryo medium. The tank was mounted on a shaker and oscillated around its central axis at a frequency of 1 Hz to prevent the larvae from entering a sleep state. Continuous light exposure was provided throughout the 24 h deprivation period. Paramecia were supplied twice daily to prevent starvation. The medium was replaced daily to maintain optimal water quality. All experiments were conducted during the dark phase (22:00 or later).

### Auditory stimuli

Auditory stimuli were integrated into the all-optical system. Binaural sound stimuli (350 Hz, 500 ms duration, 100 dB intensity) were delivered at 60 s intervals. The experimental protocol consisted of an initial baseline period of 10–20 minutes to determine the baseline locomotor speed of each zebrafish, followed by approximately 1 hour of auditory stimulation. During the auditory stimulation period, optogenetic activation was applied using a 588 nm laser in multiple epochs, each lasting 5–10 minutes, with an inter-stimulation interval of at least 10 minutes. Zebrafish behavior was recorded at 50 fps using an infrared camera (Basler acA2000-165kmNIR), and locomotor speed was calculated from the tracked positional data.

For drug-treatment experiments, the protocol consisted of four sequential phases. First, a 20-minute baseline period was recorded. This was followed by a 30-minute auditory stimulation. Subsequently, all E2 embryo medium was removed and replaced with E2 medium containing 1 mM mepyramine. After a 30-minute incubation period to allow the drug to take effect, auditory stimulation was applied again for an additional 30 minutes.

In freely swimming zebrafish, an escape response was defined as an increase in swimming speed exceeding 50% of the baseline average within 1 s following stimulus onset. In head-fixed zebrafish, a DeepLabCut model was trained to detect six key points along the tail to extract locomotor kinematics. Bouts were identified based on the cumulative tail curvature over 0.5 s sliding window, after removing slow trends using a moving average filter with a 9 s window. Bouts occurring within 1 s of auditory stimulus onset were considered stimulus-evoked.

### dPCA

dPCA was performed using the MATLAB implementation provided by the Machens laboratory (https://github.com/machenslab/dPCA). Head fixation ensured stable whole-brain recording and reliable dimensionality reduction. Prior to analysis, whole-brain neural activity traces were z-score normalized across time. To reduce the influence of sampling noise and motion artifacts, brain regions with volumes smaller than 50 voxels were excluded.

For the identification of the DRN 5-HT activation–related neural subspace, all data were first registered to the standard brain, after which the DRN region itself was removed to prevent direct optogenetic activation effects from dominating the observed activity differences. Neural activity was segmented into trials based on the presence or absence of optogenetic stimulation, without regard to the behavioral state of the fish or the presence of auditory stimuli. Each trial had a fixed duration of 11 s. dPCA was then applied to these trial-structured data. After applying dPCA, we found that in *Tg(tph2:ChrimsonR)* zebrafish, the first demixed principal component (dPC1) explained more than 50% of the total variance, whereas the remaining dPCs accounted for substantially smaller proportions. Therefore, all subsequent analyses were based on dPC1, which was defined as the DRN activation–related subspace. For visualization, dPC1 score for each fish was normalized by its maximum score value and the traces were aligned to DRN optogenetic stimulation onset. Brain regions with high absolute weights on dPC1 were selected, and their spatial distributions were visualized. A linear regression model was used to quantify the fraction of variance in these regions that could be explained by motor behavior (*R*^2^; Methods). To assess the specificity of this relationship, an equal number of neurons was randomly sampled from the remaining whole-brain population (excluding previously removed regions), and their *R*^2^ values were calculated and statistically compared with those of the dPC1-weighted regions.

To identify sound-evoked and motor-correlated neural subspaces, neural activity was categorized into four trial types according to the presence or absence of auditory stimulation and locomotor behavior. Trials containing optogenetic stimulation were excluded from this analysis to avoid confounding effects. Neural activity was temporally aligned either to the onset of auditory stimuli or to the initiation of locomotor bouts. dPCA was then applied using auditory stimulation or movement as the decoding variable to extract sound-related and motor-related neural components. Because the first demixed principal components (dPC1-sound and dPC1-motor) consistently accounted for a substantially larger fraction of the variance than higher-order components, whole-brain activity was projected onto these components.

### Similarity analysis

To quantify the similarity of population neural responses in the sound-evoked subspace across trials under different optogenetic and drug-induced conditions, we constructed trial-by-trial similarity matrices using cosine similarity. Neural activity was z-score normalized across time for each neuron prior to analysis. Trials were defined as the time of auditory stimulus onset and the following 2 s. For each trial, neural activity within the corresponding time window was extracted. Activity for each neuron was then averaged over time to obtain a neurons × 1 vector representing the population response. Population response vectors from all trials were concatenated to form a neurons × trials matrix. Pairwise similarity between trials was computed using cosine similarity across neurons.

### Subspace angle analysis

To assess the alignment between the DRN activation–related neural subspace and either the motor-related or sound-evoked subspaces, we quantified subspace similarity using principal angles. For each subspace, the first two dimensions were retained to define the corresponding low-dimensional representation. Orthonormal bases for the DRN activation, motor-related, and sound-evoked subspaces were obtained using the MATLAB function `orth`. Principal angles between each subspace and the DRN activation subspace were then computed using the `subspace` function.

To evaluate the statistical significance of the observed subspace alignment, we performed a nonparametric permutation test. To generate a null distribution, 1,000 random subspaces were constructed by randomly selecting time points from the spontaneous neural activity matrix and orthonormalizing the resulting activity vectors using `orth`. For each random subspace, principal angles relative to the DRN activation subspace were computed using the same procedure, yielding a null distribution of mean principal angles under the hypothesis of random alignment. The observed mean principal angle between the DRN subspace and either the motor-related or sound-related subspace was then compared against the null distribution. Two-tailed empirical p-values were calculated as the proportion of permuted means that were equal to or more extreme than the observed value.

### Linear regression

We constructed a linear regression model based on kinematic features extracted from tail motion to quantify how tail movements explain whole-brain neural activity in larval zebrafish. Swimming direction and distance were derived from tail movements, with distance defined as the area swept by the tail during a bout and direction defined as the cumulative tail bending angle within a single bout. Based on these two parameters, a polar coordinate system was constructed, and swimming bouts were classified into 12 discrete types.

For each bout type, a corresponding binary regressor was generated, which took a value of 1 during the occurrence of that bout and 0 otherwise. Each regressor was then convolved with the experimentally measured jGCaMP8s calcium response kernel. The resulting regressors formed the matrix *X*.For each neuron, the linear regression model was defined as

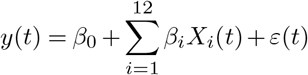

where *y*(*t*) denotes the neural activity time series, *X*_*i*_(*t*) represents the convolved regressor for the *i*-th bout type, *β*_*i*_ is the corresponding regression coefficient, *β*_0_ is the intercept, and *ε*(*t*) denotes the residual error.

Model fitting was performed using the MATLAB function `fitlm` with ordinary least squares estimation. The coefficient of determination (*R*^2^) was calculated for each neuron to quantify the proportion of variance in neural activity explained by motor behavior.

For each fish, to avoid confounding effects, only tail movement data recorded during the baseline period without optogenetic stimulation or auditory stimulation were used to construct the model. Fish whose baseline tail movements were insufficient to cover all 12 bout types were excluded from further analysis. Motor-related neurons were then identified based on both the proportion of variance in neural activity explained by movement (*R*^2^) and their weights in the motor-related subspace. To quantify the effect of optogenetic stimulation on motor-related neurons, the model fitted during the baseline period was used to predict their neural activity during both the control (no stimulation) and DRN 5-HT activation periods.

### Hyperbolic and Euclidean Multidimensional Scaling

We analyzed the geometry of motor population activity using multidimensional scaling (MDS) on pairwise neural correlation distance matrices. Pairwise distances between motor-correlated neurons were computed from the Pearson correlation matrix *C* as

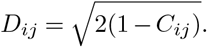

This is the chord distance on the unit hypersphere of *z*-scored activity vectors and is a proper metric (*D*_*ij*_ = *D*_*ji*_, *D*_*ii*_ = 0, triangle inequality satisfied), which is required for the validity of the Bayesian MDS embedding and the Bayesian Information Criterion (BIC) based dimensionality selection. Distances are bounded in [0, 2], with *D*_*ij*_ = 0 for perfectly correlated, 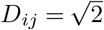 for uncorrelated, and *D*_*ij*_ = 2 for perfectly anticorrelated neurons.

Our main analysis used Bayesian hyperbolic MDS (HMDS) to embed the data in a Poincaré ball, following Praturu et al. (58). Using the `metric_HMDS` Python library, we jointly optimized the embedded coordinates and a global curvature parameter *λ* (constant negative curvature *K* = − *λ*^2^) so that hyperbolic distances *δ*_*ij*_ best matched the neural distances *D*_*ij*_:

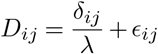

The embedding dimension (*d* = 6) was chosen by minimizing the Bayesian Information Criterion (BIC) over candidate dimensions, defined as

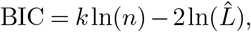

where 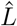 is the maximized model likelihood, *k* is the number of free parameters, and *n* is the number of observations. For a distance matrix of *N* points, *n* = *N* (*N* − 1)*/*2. For the HMDS model with embedding dimension *d*, the effective number of parameters is

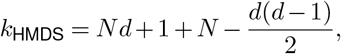

accounting for *N* points in *d* dimensions, one global curvature *λ, N* point-specific uncertainty parameters, and adjusting for the rotational symmetry of hyperbolic space. We found the optimal embedding *d* by minimizing BIC (Fig. S7B). For comparison, we applied classical Euclidean MDS (`scikit-learn, dissimilarity=‘precomputed’, metric=True`) with the same dimension. Its BIC used *N × d* coordinate parameters and one global variance parameter.

To estimate intrinsic variability in pairwise neural correlation distances from non-stationary brain dynamics, we performed a time-window–based uncertainty analysis. Each recording was divided into non-overlapping 60 s windows. For each window, we computed population activity vectors and the corresponding pairwise correlation distance matrix. We then calculated, for each unique pair, the variance of its distance across windows, yielding a distribution of temporal stability. This produced a variance matrix Σ_*D*_ with the same dimensions as the averaged distance matrix *D*, where Σ_*D*_(*i, j*) is the empirical temporal variance of the distance between neurons *i* and *j*. These variance estimates were incorporated into both HMDS and Euclidean MDS by weighting each pairwise distance in the likelihood function inversely by its variance.

We visualized the embeddings to interpret the geometry. For dimensions > 3, we projected the coordinates into 3D using PCA and displayed them within the Poincaré ball. Embedding quality was evaluated with Shepard diagrams (Fig. S7C), plotting original distances against embedded distances. A perfect embedding would place all points on the *y* = *x* diagonal.

### Eigenvector-randomization surrogate

To test whether the fitted hyperbolic curvature reflects genuine geometric structure in the motor population rather than spectral properties of the shared variance alone, we generated surrogate datasets that preserve the eigenvalue spectrum of the covariance matrix while destroying the arrangement of which neurons participate in which mode.Specifically, we computed the covariance matrix Σ of the activity traces across motor-correlated neurons and

eigendecomposed it as Σ = *U* Λ *U*^⊤^. We then replaced the eigenvector matrix *U* with a random orthonormal basis *U*^*′*^, drawn uniformly from *O*(*N*) via QR decomposition of a random Gaussian matrix, and reconstructed a surrogate covariance matrix Σ*′* = *U*^*′*^ Λ *U*^*′*⊤^. The surrogate covariance was converted to a correlation matrix 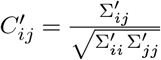 from which pairwise distances were computed. We applied the full HMDS pipeline to *C*^*′*^ under identical settings and compared the fitted curvature *λ* and the Shepard diagram to those from the original data (Fig. S9B).

### Cross-animal analysis of curvature and behavior

For each fish, we computed bout frequency separately for laser-on and laser-off periods as swim bouts per second. Bouts were detected from the tracked tail trace using the cumulative tail-curvature criterion described above (“Bout detection”). Each correlation distance matrix was normalized by a scaling factor such that the maximum distance *D*_*ij*_ was equal to 2. Curvature was fit independently for each fish and condition using the HMDS procedure above. To obtain an estimate of fit uncertainty for *λ*, we repeated the HMDS optimization multiple times per fish and per condition from different random initializations. The reported *λ* values are mean *±* SD across these repeats. Fit uncertainty was systematically larger during DRN activation than during baseline, consistent with a more strongly warped embedding being harder for the optimizer to localize precisely. We quantified associations between bout frequency and fitted curvature *λ* with Spearman rank correlation *ρ*, and obtained two-sided *p*-values by exact permutation: for each comparison, we evaluated all 7! = 5040 permutations of *λ* across fish, computed *ρ* for each, and took the fraction with |*ρ*_perm_ | ≥ | *ρ*_observed_ | as the exact *p*-value. This avoids the asymptotic assumptions of the standard Spearman test, which are unreliable at *n* = 7. Pearson correlations and their parametric *p*-values are reported for comparison.

### Statistical analysis

Wilcoxon matched-pairs signed rank test and Mann–Whitney U test were used and data are expressed as the mean *±* min-max in Fig. 2C. Mann–Whitney U test was used in Fig. 1D. Mann–Whitney U test was used and data are expressed as the mean *±* min-max in Fig. 2D and Fig. 3D. Wilcoxon matched-pairs signed rank test was used in Fig. 4B. P-values obtained via a nonparametric permutation test in Fig. 4F. Data are expressed as the mean *±* SD in Fig. 1D, Fig. 2B, Fig. 5B, Fig. 5C.

## Supplementary figures (S1)

**Figure S1.**
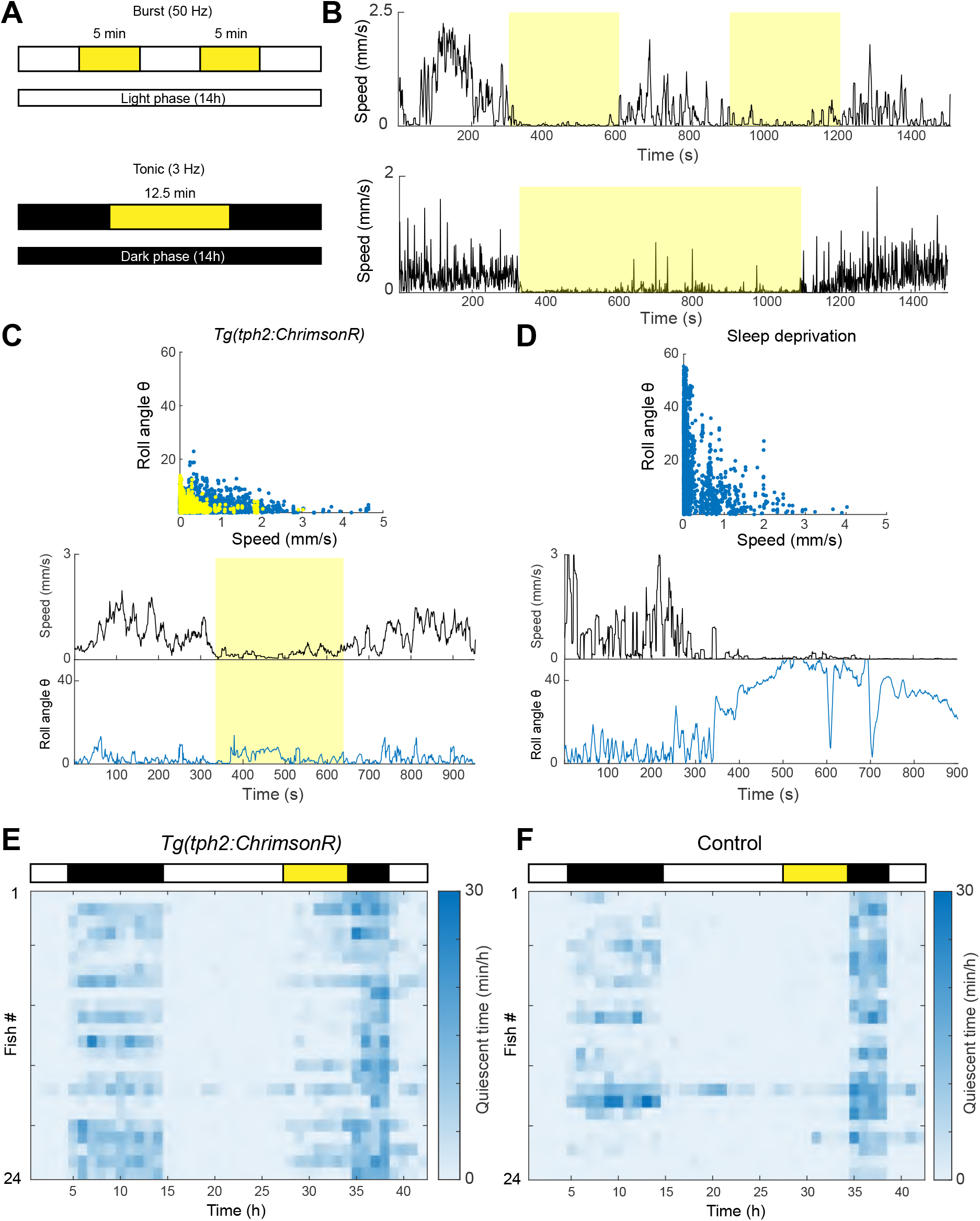
DRN 5-HT activation suppresses locomotion and induces a quiescent state distinct from sleep. **A**. Schematic of burst and tonic optogenetic stimulation paradigms. **B**. Top: locomotor speed of a zebrafish during burst stimulation. Bottom: locomotor speed of a zebrafish during tonic stimulation. **C**. Relationship between body roll angle and swimming speed in Tph2:ChrimsonR zebrafish. Each point represents one second; yellow points indicate periods of DRN 5-HT activation. **D**. Relationship between body roll angle and swimming speed in sleep-deprived zebrafish. **E**. Quiescence per hour in Tph2:ChrimsonR zebrafish. **F**. Quiescence per hour in control zebrafish.

**Figure S2.**
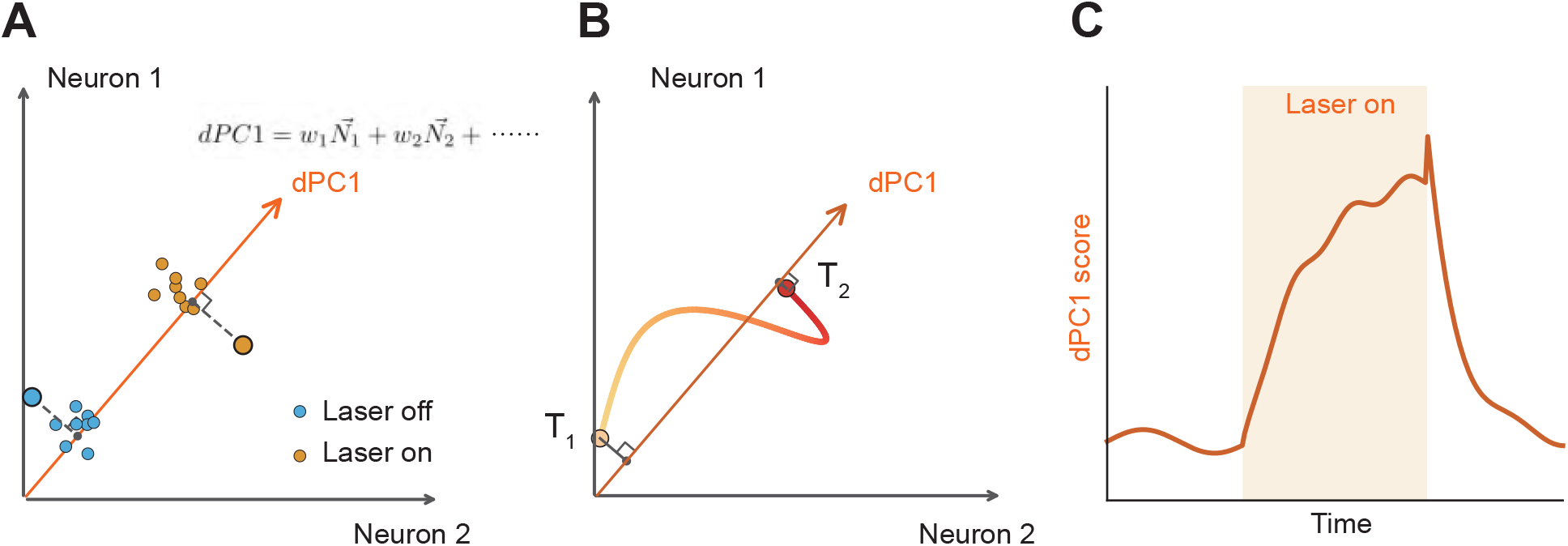
Geometric interpretation of dPCA. **A**. Neural activity at each time point is a point in an N-dimensional space defined by the recorded neurons (illustrated for two neurons). dPCA finds a direction (dPC1, orange arrow) that best separates conditions — here, DRN-activated (orange) versus control (blue). Dashed lines show orthogonal projections of individual time points onto the dPC1 axis. The weights defining dPC1 as a linear combination of the neural axes are shown below. **B**. Over time, the population state traces a trajectory through this space (color gradient from light to dark indicates progression from *T*_1_ to *T*_2_). Projecting this trajectory onto dPC1 (dashed lines with right-angle marks) gives a one-dimensional time course of the activation-related component of population activity. **C**. The resulting dPC1 score, plotted as a time series with the laser-on period shaded, is the quantity shown in Fig. 3B and reflects how strongly the population is driven along the activation-related direction at each moment.

**Figure S3.**
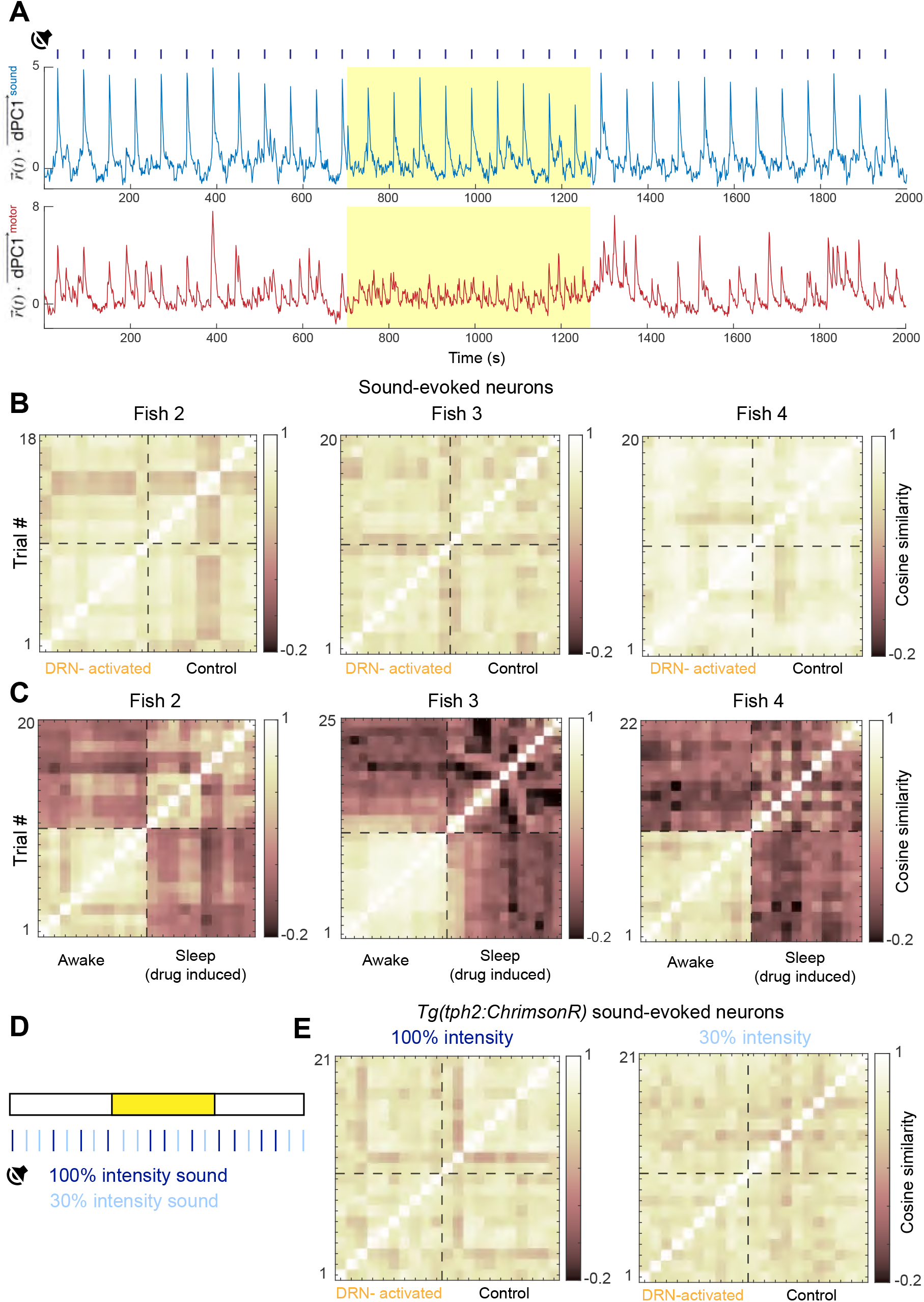
DRN 5-HT activation did not alter neural dynamics within the sound-evoked subspace. **A**. Temporal evolution of whole-brain neural activity projected onto the sound-evoked subspace and the motor-correlated subspace in an example Tg(tph2:ChrimsonR) zebrafish. Yellow shading indicates periods of optogenetic stimulation. **B**. Similarity matrices of sound-evoked neuronal population responses during DRN 5-HT activation and control periods for fish 2–4 (fish 1 shown in Fig. 4E). **C**. Same analysis as in panel B, but comparing the awake state with the drug-induced sleep state. **D**. Schematic of auditory stimulation paradigms with different sound intensities. **E**. Similarity matrices of sound-evoked neuronal population responses for strong (left) and weak (right) sound stimuli during DRN 5-HT activation and control periods.

**Figure S4.**
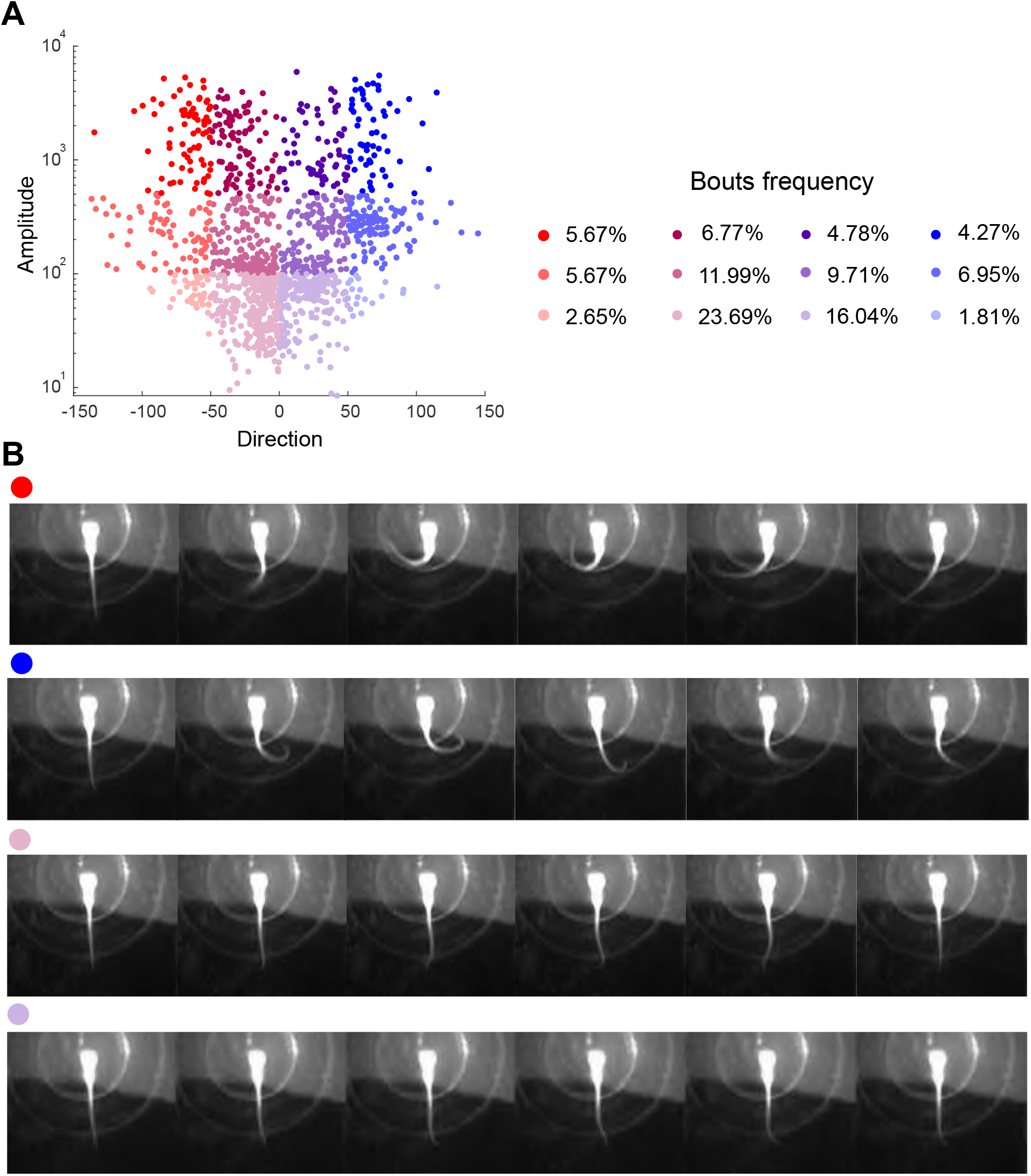
Distribution of bouts in the direction–amplitude space. **A**.Left: Distribution of all detected bouts projected onto the direction–amplitude space. Each bout is color-coded according to the classification scheme used in the main text (n = 7 fish, 1,493 bouts). Right: Frequency distribution of each bout type. **B**. Representative examples of distinct bout types, illustrating both large- and small-amplitude tail deflections. Colors are consistent with those used in the main figures.

**Figure S5.**
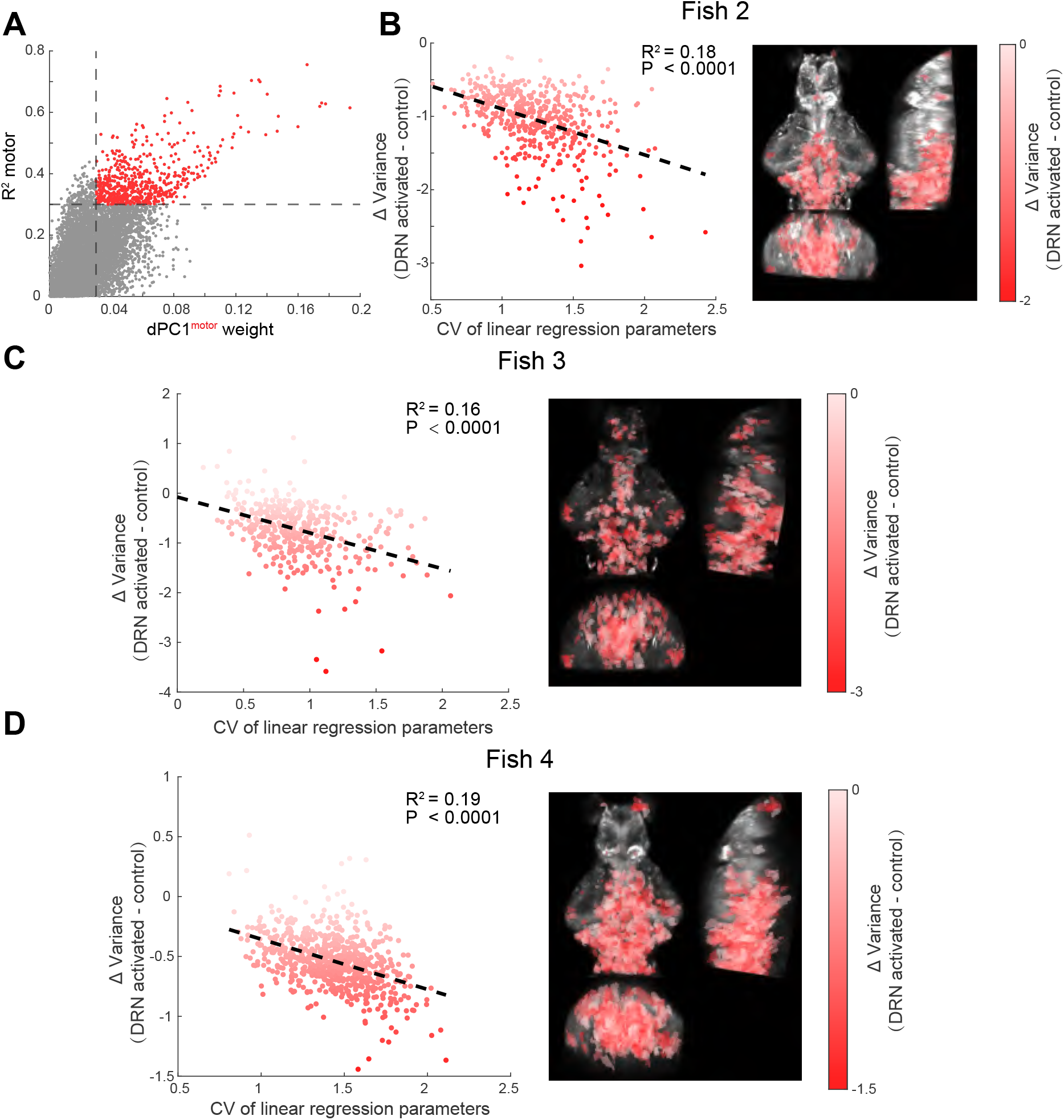
DRN 5-HT activation exerts graded suppression on the motor subspace. **A**. Relationship between *dP C*1^*motor*^ weights and *R*^2^. Red points indicate the motor-correlated neurons included in the analysis. **B-D**. Relationship between DRN 5-HT activation–induced modulation of neural activity and the coefficient of variation (CV) of regression coefficients in motor-correlated neurons for fish 2–4 (fish 1 shown in Fig. 5D–E).

**Figure S6.**
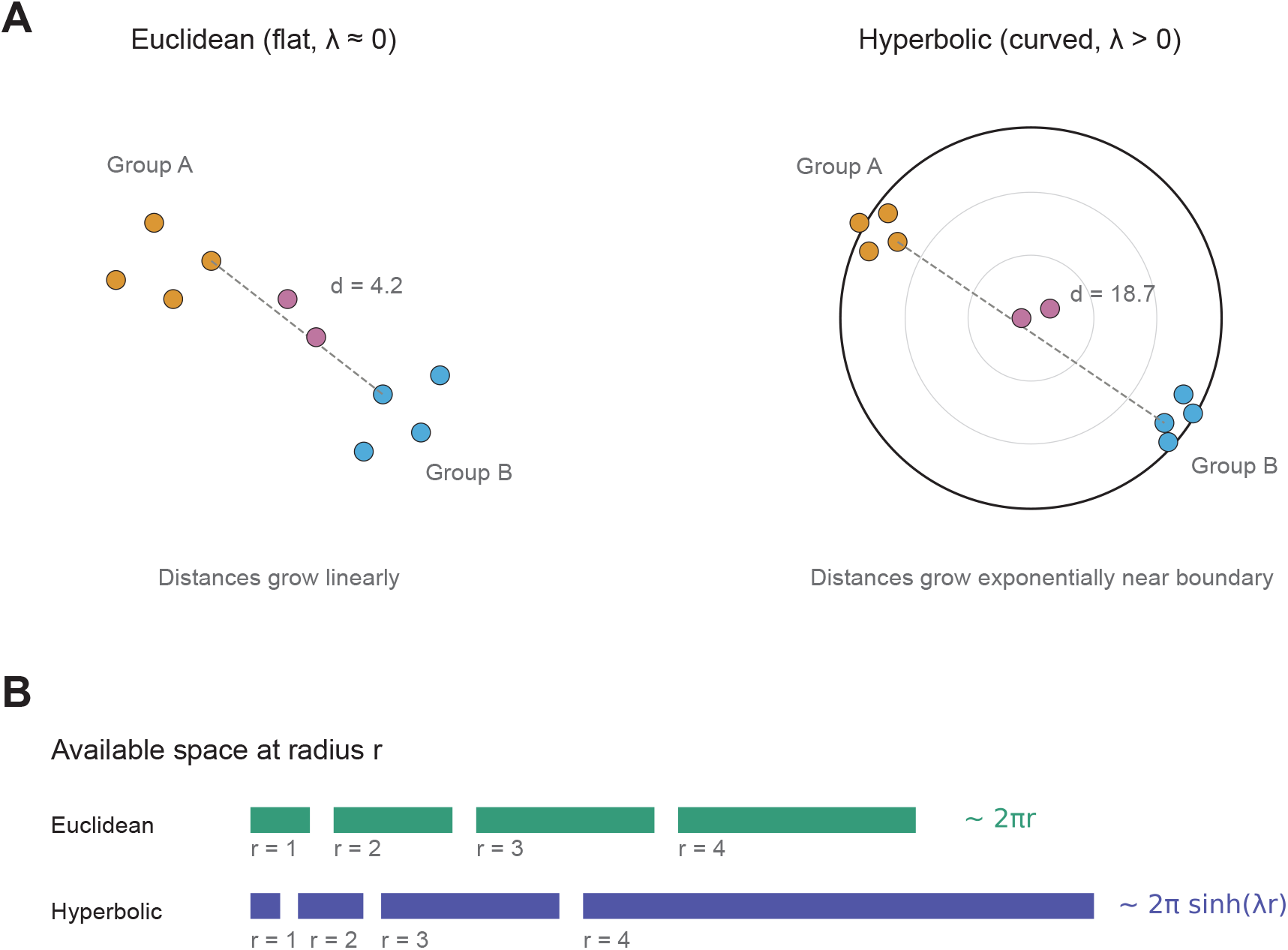
Intuition for hyperbolic versus Euclidean embedding. The same neuron population (colored by functional group) embedded in flat Euclidean space (left) and in a Poincaré disk representation of hyperbolic space (right). In Euclidean geometry, distances between groups grow linearly, so separation is moderate. In hyperbolic geometry, space near the disk boundary expands exponentially, allowing functionally distinct groups (orange, blue) to be far more separated than in an equally dimensional Euclidean space, even though they appear visually close in the disk. Dashed lines show the distance between the same two groups in each geometry. Pink points near the center lie at moderate distance from both groups. Bottom inset: available space (circumference) at radius r grows as ∼ 2*πr* in Euclidean geometry but as ∼ 2*π* sinh *λr* in hyperbolic geometry, where *λ* is the curvature parameter. This exponential expansion lets hyperbolic embeddings accommodate populations with strongly segregated functional subsets.

**Figure S7.**
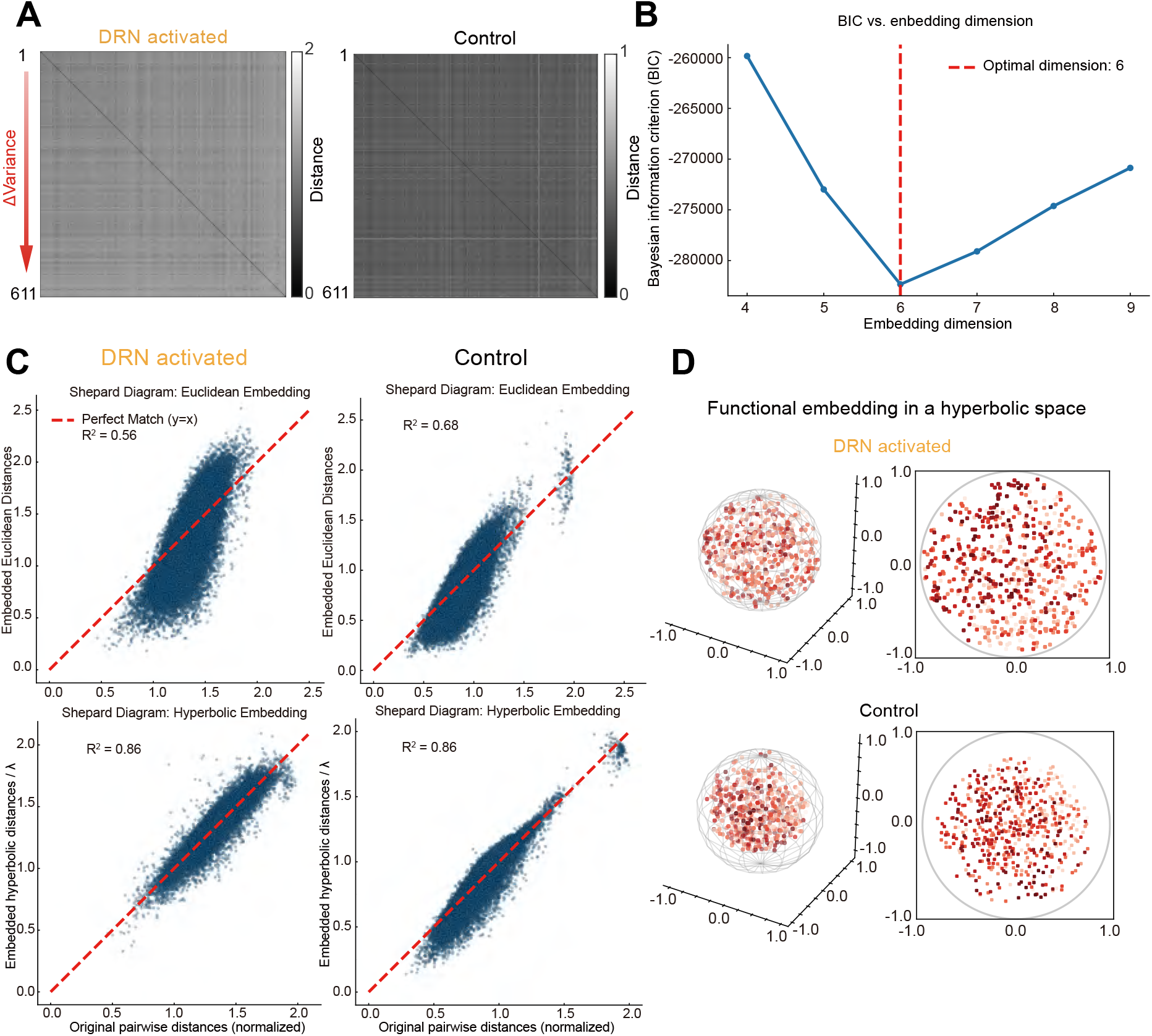
DRN 5-HT activation diversifies motor network activity. **A**. Pairwise distance matrices of motor-correlated neurons during DRN activation (left) and control (right) periods (fish 1). Neural activity magnitude was quantified by the variance. Neurons were ranked by changes in variance between DRN 5-HT activation and control periods. **B**. Embedding dimension as a function of BIC, see Methods. **C**. Shepard diagram of embedded vs. original pairwise distances for fish 1. Original distance matrices were scaled so the maximum distance is 2. Left: DRN activation (laser on); Right: baseline (laser off). Top: Euclidean embedding; bottom: hyperbolic embedding. As indicated by R^2^, hyperbolic embedding outperforms Euclidean embedding under all conditions, consistently across all fish. Embedding dimension *d* = 6. **D**. Hyperbolic multidimensional scaling (HMDS) of neural correlation distances in a 3D Poincaré ball (left) and 2D projection (right), see Methods. Top: functional embedding during DRN 5-HT activation; bottom: control. In the Poincaré ball, distances diverge near the boundary. During DRN activation, two neuronal ensembles segregated to opposite poles (light and dark red, as in Fig. 5E), indicating increased functional diversity, whereas at baseline the embedding was more compact.

**Figure S8.**
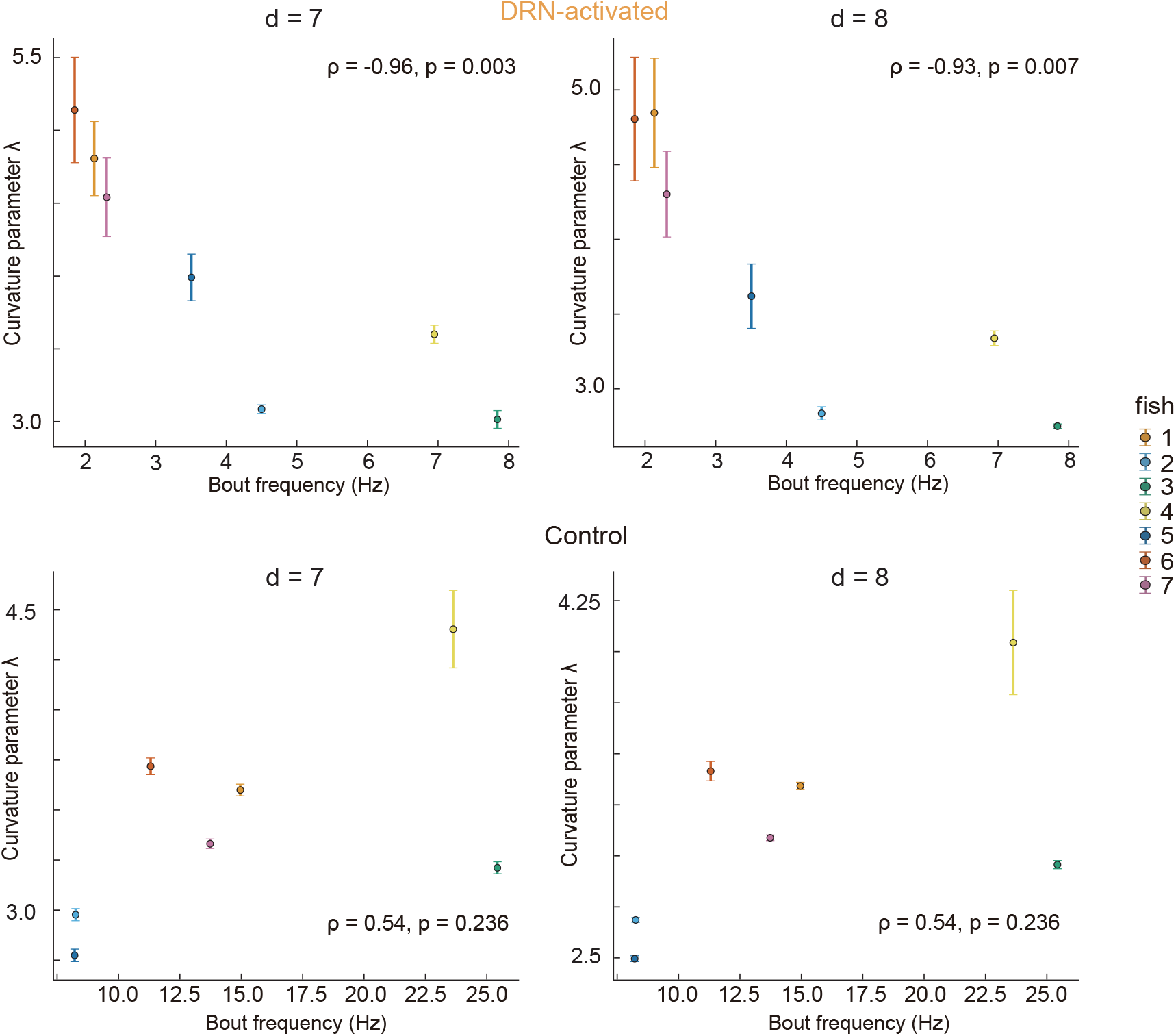
Curvature–behavior relationship is robust across embedding dimensions. Same analysis as Fig. 5F but with embedding dimensions d = 7 (left column) and d = 8 (right column). Top row: during DRN activation, *λ* correlates negatively with bout frequency at both d = 7 (Spearman *ρ* = 0.96, p = 0.003) and d = 8 (Spearman *ρ* = 0.93, p = 0.007, exact permutation tests). Bottom row: no significant relationship during baseline in either dimension. Each point is one zebrafish (n = 7), colored by individual; error bars, *±* 1 SD from the HMDS fit. All p-values computed by exact permutation (5040 permutations).

**Figure S9.**
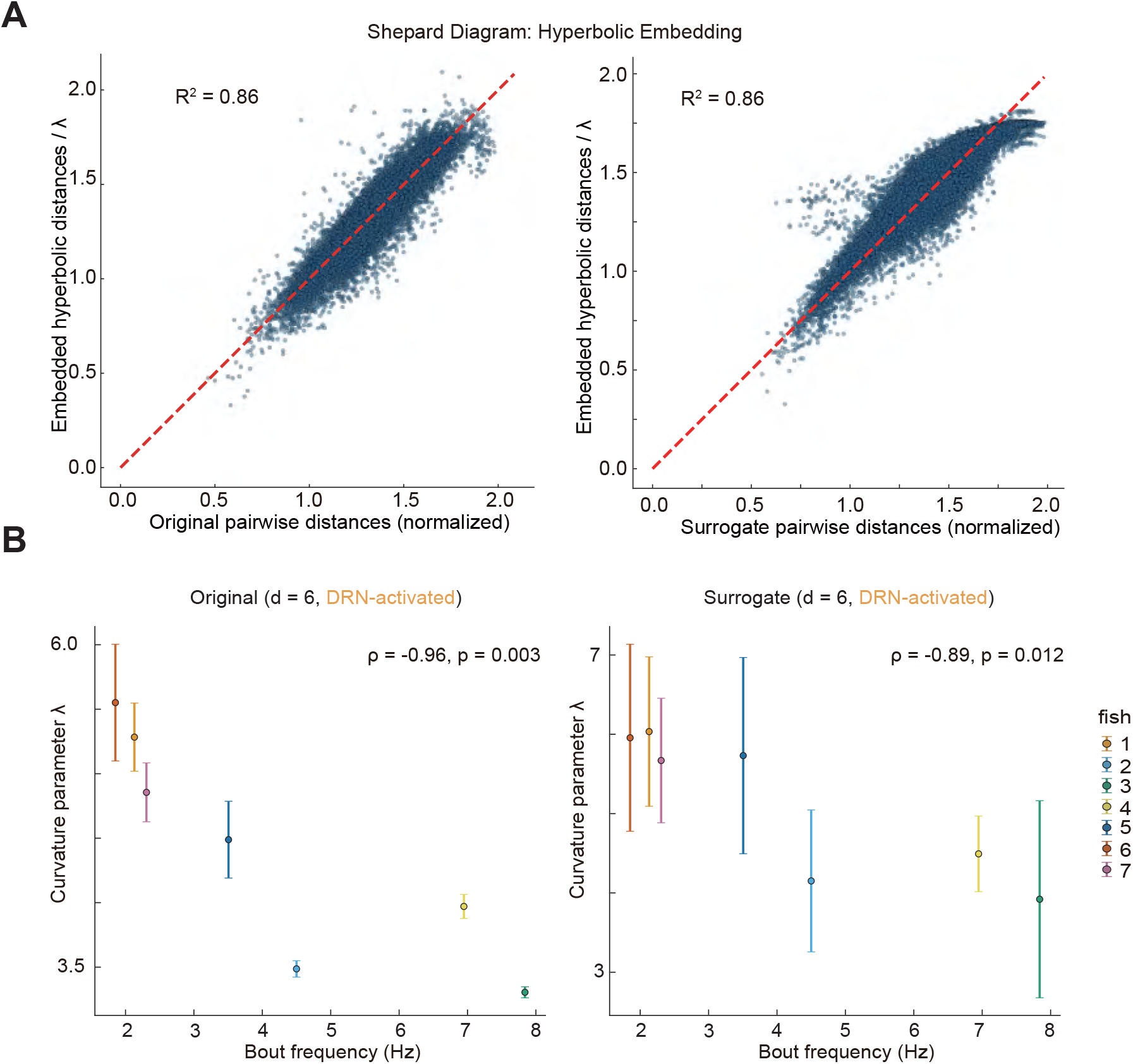
Eigenvector-randomized surrogate distorts pairwise correlation distances. **A**. Shepard diagrams for hyperbolic embedding of original (left) and surrogate data (right). Left, original motor-population data (fish 1) during DRN activation. Points track the diagonal with approximately symmetric scatter. Right, covariance-eigenvector-randomized surrogate. The surrogate was generated by replacing the eigenvectors of the covariance matrix with a random orthonormal basis while preserving the eigenvalue spectrum (Methods). The cloud systematically deviates below the diagonal at large distances, indicating that the surrogate embedding compresses distances between dissimilar neurons. This distortion is absent in the original data, confirming that the quality of the hyperbolic fit depends on the specific geometric arrangement of neurons in correlation space, not solely on the eigenvalue spectrum of the covariance matrix. Shepard diagrams for other fish exhibit qualitatively similar behaviors. **B**. Curvature parameter *λ* vs. laser-on bout frequency for original data (left) and surrogate data (right), both at d = 6. Each point is one zebrafish (n = 7); error bars, *±* 1 SD across repeated HMDS fits. The surrogate yields substantially larger fit uncertainty, indicating that the optimizer cannot converge on a stable embedding in the absence of a specific geometric structure. The tight curvature–behavior correlation observed in the original data is also weakened in the surrogate.

